# Modelling the interaction between silver(I) ion and proteins with the 12-6 Lennard-Jones potential: a bottom-up parameterization approach

**DOI:** 10.1101/2024.11.28.625676

**Authors:** Luca Manciocchi, Alexandre Bianchi, Valérie Mazan, Mark Potapov, Katharina M. Fromm, Martin Spichty

## Abstract

Silver(I) ions and organometallic complexes thereof are well established antimicrobial agents. They have been employed in medical applications for centuries. It is also known that some bacteria can resist to silver(I) treatments through an efflux mechanism. However, the exact mechanism of action remains unclear. All-atom force-field simulations can provide valuable structural and thermodynamic insights on the molecular processes of the underlying mechanism. Lennard-Jones parameters of silver(I) have been available for quite some time; their applicability to describe properly the binding properties (affinity, binding distance) between silver(I) and peptide-based binding motifs is, however, still an open question. Here, we demonstrate that the standard 12-6 Lennard-Jones parameters (previously developed to describe the hydration free energy with the TIP3P water model) significantly underestimate the interaction strength between silver(I) and both methionine and histidine. These are two key amino-acid residues in silver(I)-binding motifs of proteins involved in the efflux process. Using free-energy calculations, we calibrated non-bonded fix (NBFIX) parameters for the CHARMM36m force field to reproduce the experimental binding constant between amino acid sidechain fragments and silver(I) ions. We then successfully validated the new parameters on a set of small silver-binding peptides with experimentally known binding constants. In addition, we could monitor how silver(I) ions increase the α-helical content of the LP1 oligopeptide, in agreement with previously reported Circular Dichroism (CD) experiments. Future improvements are outlined. The implementation of these new parameters is straightforward in all simulation packages that can use the CHARMM36m force field. It sets the stage for the modeling community to study more complex silver(I)-binding processes such as the interaction with silver(I)-binding-transporter proteins.

## 1. Introduction

The antimicrobial properties of silver(I) ions are well known and applied in various medical solutions.^1–3^ However, some bacteria are known to resist to silver(I) cytotoxicity through an efficient efflux pump that expels the metal ion out of the cell.^4^ The efflux mechanism is based on the interplay between the small periplasmic silver(I)-transporter protein SilF and the tripartite efflux complex SilCBA; the latter functions as a chemiosmotic metal ion/proton antiporter. In addition, the disordered protein SilE acts as molecular sponge^5^ to absorb silver(I) ions in the periplasm; probably to regulate the metal ion concentration.^6^ However, the precise role of SilE remains unclear.

Particularly important for the binding of silver(I) ions to SilE and other proteins of the efflux machinery are the amino acids histidine (H) and methionine (M); the motifs HXXM and MXXH are privileged binding motifs for silver(I) ions.^7,8^ Their impact, however, on the dynamic structure of the peptide and the binding affinity is not fully understood yet. An interesting observation is also that silver(I) binding to SilE and non-structured peptides induces the formation of structured, alpha-helical motifs. The actual molecular mechanism remains unclear.^4,8,9^

All-atom molecular dynamics simulations conducted on the nanosecond to microsecond timescale could provide valuable insights into these unresolved questions. Force-field parameters for silver(I) ions have been developed previously: Won parameterized mono-,di and tri-valent ions, including silver(I), to reproduce the hydration free energy using Jorgensen’s TIP3P^10^ water model.^11^ To our understanding, these parameters were unfortunately not correctly integrated into the official distribution of the CHARMM force field. The epsilon values lack the proper conversion from kilojoules to kilocalories. This issue is currently reviewed by the developers of the CHARMM force field (Alex MacKerell; personal communication, 5 December 2024). Merz and coworkers determined alternative 12-6 LJ silver(I) parameters. They aimed to reproduce either the hydration free energy (*HFE*) or the ion-oxygen distance (*IOD*) for multiple water models.^12^ A 12-6-4 LJ model was also proposed to account for polarizability. The Merz group later proposed specific 12-6-4 LJ parameters for the interaction of silver(I) (among other metal ions) with imidazole and acetate.^13,14^ While the 12-6-4 LJ potential is certainly a valuable extension it is not available in most simulation packages by default. Thus, we limit our focus in this work to the standard 12-6 Lennard-Jones potential.

We first assess whether the aforementioned 12-6 Lennard-Jones parameters can be directly applied to study the interaction of silver(I) ions with the HXXM binding motif as described by the CHARMM36m^15^ force field. We then describe the development of improved 12-6 Lennard-Jones parameters for modeling the interaction of silver(I) with histidine, methionine, and phenylalanine. We thereby employed a bottom up strategy: we first developed parameters for the silver(I) interaction with sidechain fragments for which affinity data was available from literature or that we measured by potentiometric or spectrophotometric titrations. We then validated the derived parameters against binding constants data for HXXM and MXXH motifs, as measured by NMR titrations.^7,8^ Finally, as an application of the new parameters, we tested them against structural data for the oligopeptide LP1 (also called B1b^9^), in particular the transition from an intrinsically disordered peptide to an alpha-helical structure in the presence of silver(I) ions.

## 2. Methodology

This section provides a general outline of the chosen methodology. Details of the applied computational procedures are given in the Electronic Supplementary Information (ESI).

### 2.1 Non-bonded interaction potential

In the family of CHARMM force fields the non-bonded interaction between a pair of atoms is given by the following potential:

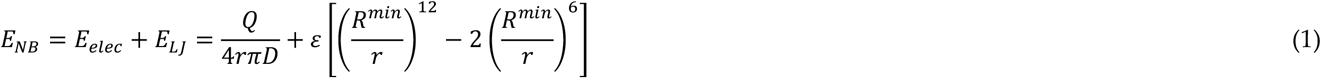

where *r* is the distance between the two atoms, *Q* is the product of their partial charges, *D* is the product of the vacuum permittivity and the relative permittivity of the medium (=1 for explicit water simulations). The parameters *ε* and *R*^*min*^ are in general derived from the Lorentz-Berthelot rule:^16,17^

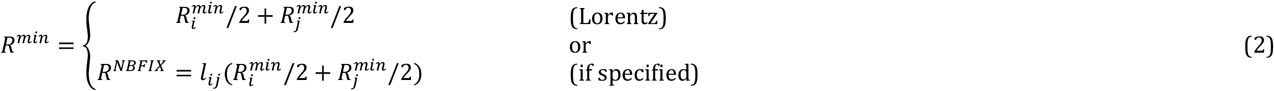

and

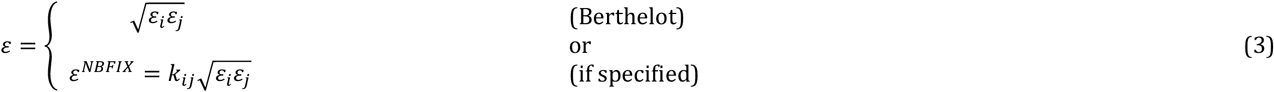

where the subscript indicates an atom-type specific parameter of the *i*-th and *j*-th atom. Beside this general combination rule, it is also possible with the NBFIX feature of CHARMM to directly specify *ε*^*NBFIX*^ and *R*^*NBFIX*^ for an interaction between two specific atom types (instead of using the Lorentz-Berthelot rules). Alternatively, *ε*^*NBFIX*^ and *R*^*NBFIX*^ can expressed also in terms of pairwise correction coefficients *k*_*ij*_ and *l*_*ij*_, respectively.^18^ It allows to selectively modify the strength and distance for a particular interaction without affecting the other interactions. In what follows we refer to the pair *ε*^*NBFIX*^ and *R*^*NBFIX*^ as NBFIX parameters.

### 2.2 Preliminary evaluations

As an initial test, Won’s parameter^11^ set and Merz’ *HFE* and *IOD* parameter sets^12^ for silver(I) ions were tested in combination with the CHARMM36m force field^15^ using the general Lorentz-Berthelot rule. The strong silver-binding tetrapeptide HEFM was chosen as test case because it includes four amino acid sidechains that are supposed to bind silver. As shown in the section 3.1 and in Table 1, all three sets drastically underestimated the binding affinity of HEFM and silver(I).

**Table 1.** Results for the initial MD simulations of the HEFM tetrapeptide. Three blocks of simulations of 100 ns each were performed for two box sizes and for each parameter set. Values are given with standard error of the mean from block analysis.

| Box size (Å) | Number of peptides | Number of Ag<sup>+</sup> ions | Parameter set | Affinity $\log_{10}(K_{bind,calc})$ |
| --- | --- | --- | --- | --- |
| 30 | 1 | 1 | Won<sup>a</sup> | 0.8 ± 0.1 |
|  |  |  | HFE<sup>b</sup> | 1.0 ± 0.1 |
|  |  |  | IOD<sup>c</sup> | 0.5 ± 0.1 |
| 50 | 1 | 1 | HFE<sup>b</sup> | 0.8 ± 0.2 |
|  |  |  | IOD<sup>c</sup> | 0.6 ± 0.1 |

<sup>a</sup> Obtained from interpolation of hydration free energy maps.<sup>11</sup> <sup>b</sup> Optimized to reproduce the hydration free energy.<sup>12</sup> <sup>c</sup> Optimized to reproduce the metal ion-water oxygen distance.<sup>12</sup>

An intuitive strategy to analyze and eventually correct the underestimated affinity is to adopt a bottom-up approach by studying the interaction of each amino acid sidechain with silver(I) separately.^19^ We therefore represented each amino acid as a sidechain fragment by removing the backbone and converting the C^β^ and H^β^ atoms into a methyl group (Figure 1); for example, ethylmethylsulfide was chosen as sidechain analogue for methionine. For histidine we converted the C^α^ and H^α^ atoms due to technical reasons (see ESI) and therefore used 4-ethylimidazole as sidechain fragment. By applying the *HFE* parameter set of Merz,^12^ the binding constant was then determined for each fragment with the following procedure:

**Figure 1.**
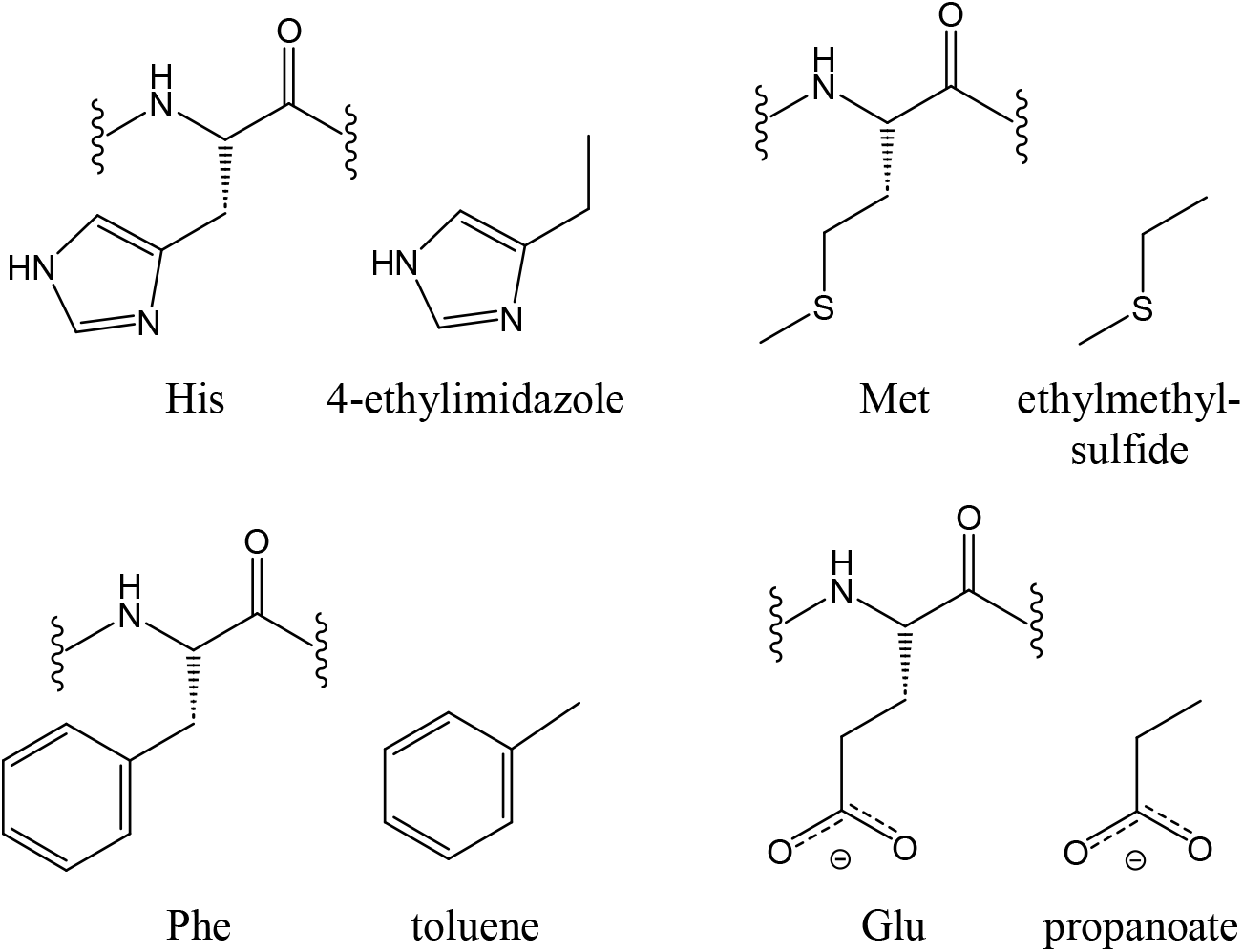
Sidechain fragments to represent histidine, methionine, phenylalanine and glutamic acid.

1. We performed MD based Umbrella Sampling (US) simulations in explicit water to sample along the reaction coordinate of binding, i.e. the distance between silver(I) ion and the coordinated atom;^20^
2. The Potential of Mean Force (PMF) was determined along the binding reaction coordinate through WHAM;^21^
3. From the PMF, the binding constant log_10_(*K*_*bind*_,_*calc*_) was calculated (and, consequently, the binding free energy Δ*G*_*bind,calc*_).^22^
4. We then compared the calculated binding constant with the experimental one from NIST database^23^ or from measured values by potentiometric & spectrophotometric titrations (see ESI), and performed a calibration if necessary.

Our initial test calculations for HEFM were carried out with CHARMM’s modified TIP3P^24^ (named cTIP3P hereafter) since it is the recommended water model for the CHARMM36m force field. The parametrizations of Merz and Won, however, were carried out with Jorgensen’s original TIP3P model (sTIP3P). To probe the influence of the two TIP3P solvation models we calculated the PMFs and binding constants of the fragments for both water models.

### 2.3. Calibration of NBFIX parameters

The main goal of this work is to derive NBFIX parameters for the CHARMM36m force field to better model the silver(I)-methionine and silver(I)-histidine interactions in particular. The modeling of histidine sidechains presents a unique challenge. Depending on the pH, the histidine sidechain can exchange between four different protonation states (Figure 2): one positively charged state with both N atoms of the imidazole ring protonated (named Hsp in CHARMM), two neutral tautomers with either the proton on the N^δ^ (Hsd) or on the N^*ε*^ (Hse), and one negatively charged state with no protons on the N atoms (imidazolate-like form). The latter is very rare in biological contexts (see PDB entry 7WAA for an example) as it requires very basic conditions (pK_a_ =12-14).^25,26^ The Hse tautomer was observed by NMR experiments and in crystal structures of small silver(I)-binding peptides and analogues thereof.^9,27,28^ In addition, quantum mechanical calculations showed that 4-methylimidazole (Hse tautomer) features a higher binding affinity than 5-methylimidazole (Hsd tautomer).^29^ Thus in this work, we only parameterized the Hse tautomer (and thus 4-ethylimidazole as sidechain representative molecule), where silver(I) binds to the nitrogen N^δ^. Note, however that under certain conditions the silver(I) ion can bind to the N^*ε*^ nitrogen, for example in SilF, where the histidine is buried inside the protein and the Hsd tautomer is stabilized through specific interactions.^6^ The possibility to include the biologically relevant tautomers through constant-pH computer simulations is discussed in the section 4.

**Figure 2.**
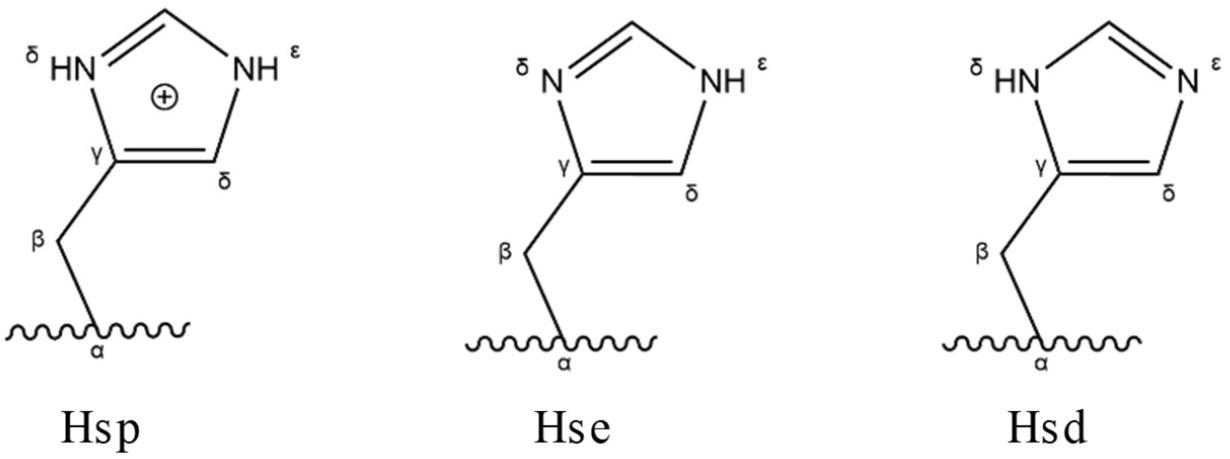
Protonation states of the histidine sidechain as defined in CHARMM force field family: Hsp (left, positive), Hse (center, neutral) and Hsd (right, neutral).

As a starting point for our optimization we selected the 12-6 LJ *HFE* parameters of Merz^12^ and determined therefrom initial NBFIX parameters for the interaction of silver(I) ion with three specific atom types of the CHARMM36m force field: NR2 (for the N^δ^ atom of histidine), S (for the sulfur atom of methionine) and CA (for the six aromatic atoms of phenylalanine). No NBFIX correction was necessary for the glutamic acid sidechain fragment. Then a calibration procedure was performed to determine the optimal NBFIX parameters that are able to correctly reproduce the experimental binding constant and binding distance of the corresponding sidechain fragments. The procedure was the following:

1. Target equilibrium distances (*d*_*eq,target*_, Table 2) between silver(I) and coordinated atoms of each fragment were taken either from the literature or determined by DFT calculations; target binding constants (log_10_(*K*_*bind*_,_*exp*_), Table 2) correspond to those of the experiments (see ESI and cited references in Table 2);
2. Using the procedure given in the section 2.2, the PMF and binding constant are calculated;
3. If the minimum of the PMF matches the target distance, we move to the next step; otherwise *R*^*NBFIX*^ is adjusted and step 2 is repeated;
4. If the calculated binding free energy is within a given error threshold of 0.5 kcal/mol, the NBFIX parameters are accepted, otherwise *ε*^*NBFIX*^ is adjusted and steps 2-3 are repeated.

**Table 2.** Optimized *ε*^*NBFIX*^ (kcal/mol) and *R*^*NBFIX*^ (Å) as well as pairwise correction coefficients *k*_*ij*_ and *l*_*ij*_ (see Eq. 1) for the interaction between silver (I) ion and specific atom types of the sidechains fragments. Results are reported for the two water models cTIP3P and sTIP3P. Calculated association constants for the corresponding sidechain fragments are reported together with the standard error of the mean from block analysis. Calculated and experimental target equilibrium distances *d*_*eq,calc*_ (corresponding to the minimum of the PMF profile) and *d*_*eq,target*_ are also reported.

| CHARMM atom type (sidechain fragment) | Water model | $\epsilon^{NBFIX}$ ($k_{ij}$) | $R^{NBFIX}$ ($l_{ij}$) | $d_{eq,calc}$<sup>a</sup> | $d_{eq,target}$<sup>b</sup> | HFE parameters<sup>c</sup> $\log_{10}(K_{bind,calc})$ | with NBFIX<sup>d</sup> $\log_{10}(K_{bind,calc})$ | Experiment $\log_{10}(K_{bind,exp})$ |
| --- | --- | --- | --- | --- | --- | --- | --- | --- |
| NR2 (4-ethylimidazole) | cTIP3P | -4.7 (116) | 2.25 (0.71) | 2.08 | 2.05-2.2<sup>e</sup> | -1.50 ± 0.02 | 3.82 ± 0.02 | 3.75<sup>f</sup>-3.85<sup>g</sup> |
|  | sTIP3P | -4.4 (109) | 2.25 (0.71) | 2.11 |  | -0.15 ± 0.01 | 3.86 ± 0.03 |  |
| S (ethylmethylsulfide) | cTIP3P | -11.9 (196) | 2.50 (0.67) | 2.55 | 2.3-2.5<sup>h</sup> | -1.89 ± 0.02 | 3.70 ± 0.02 | 3.6<sup>i</sup>-3.7<sup>j</sup> |
|  | sTIP3P | -11.85 (195) | 2.50 (0.67) | 2.54 |  | -0.29 ± 0.01 | 3.71 ± 0.03 |  |
| CA (toluene) | cTIP3P | -1.77 (74) | 2.65 (0.79) | 2.29<sup>k</sup> | 2.3<sup>k,l</sup> | -2.43 ± 0.03 | 0.43 ± 0.06 | 0.44<sup>m</sup> |
|  | sTIP3P | -1.715 (72) | 2.65 (0.79) | 2.29<sup>k</sup> |  | -2.42 ± 0.01 | 0.43 ± 0.03 |  |
| N/A.<sup>n</sup> (propanoate) | cTIP3P | N/A<sup>n</sup> | N/A<sup>n</sup> | 2.05 (2.43)<sup>o</sup> | 2.1-2.4<sup>p</sup> | 0.89 ± 0.03 | 0.89 ± 0.03 | 0.73<sup>m,q</sup> |
|  | sTIP3P | N/A<sup>n</sup> | N/A<sup>n</sup> | 2.05 (2.43)<sup>o</sup> |  | 1.25 ± 0.02 | 1.25 ± 0.02 |  |

<sup>a</sup> From the minimum of the PMF profile.
<sup>b</sup> Target equilibrium distance (see text and references below).
<sup>c</sup> Merz and coworkers.<sup>12</sup>
<sup>d</sup> Merz and coworkers<sup>12</sup> with NBFIX corrections; this work.
<sup>e</sup> Literature values.<sup>27,28</sup>
<sup>g</sup> Measured for 4(5)-ethylimidazole; this work (see ESI).
<sup>h</sup> Literature values.<sup>28,31</sup>
<sup>i</sup> Measured for ethylmethylsulfide; this work (see ESI).
<sup>j</sup> Measured for dimethylsulfide.<sup>32</sup>
<sup>k</sup> Refers to the distance from the center of the aromatic ring.
<sup>l</sup> From DFT calculations (see ESI).
<sup>m</sup> NIST database.<sup>23</sup>
<sup>n</sup> Not applicable because no NBFIX corrections are necessary (i.e. Lorentz-Berthelot rule applies)

While the chosen procedure to adjust *R*^*NBFIX*^ (step 3) and *ε*^*NBFIX*^ (step 4) should not change the outcome of the fitting process, it can impact, however, the efficiency. As a general & solid recipe we propose the following adjustment series:

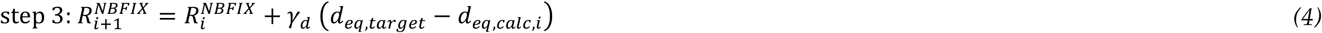

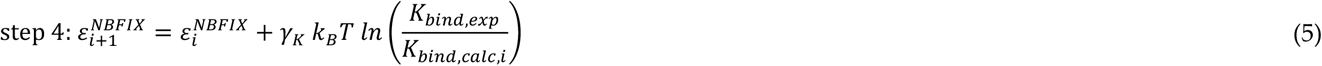

where the index *i* indicates the iteration number, and *k*_*B*_ is the Boltzmann constant and *T* the temperature. The factors *γ*_d_ and *γ*_K_ determine the step size; we typically used values between 0.5 and 1.5. The convergence can be improved by employing other root-finding techniques such as the bisection method. And even more efficient optimization strategies could make use of free-energy methods, for example, by applying multi-state acceptance ration estimators.^30^

### 2.4. Validation on HXXM and MXXH silver-binding motifs and LP1 oligopeptide

The goal of this step was to check the efficacy of the NBFIX parameters when transferred to a larger system. The validation involved calculating two different properties:

- The binding constant of four silver-binding tetrapeptides (HEFM, MNEH, HAAM and MAAH);
- The *α*-helical content in a longer peptide, LP1 (sequence: AHQKMVESHQRMMG), which was observed to form an *α*-helical structure upon addition of silver(I) ions.^9^

In both cases, potential-scaled molecular dynamics was employed by scaling down *ε* to speed up the kinetics of the dissociation process (and adjusting *R*^*NBFIX*^ so as to have the same equilibrium distance). Hamiltonian reweighting was then used to retrieve the unbiased binding constant (ESI).

To evaluate the ability of silver(I) to favor the formation of *α*-helix in LP1, two simulations were performed in parallel. In one the LP1 peptide was simulated alone in solution, whereas in the other it was simulated in the presence of one silver(I) ion. After that, the α-helical content was monitored (see ESI) and compared for the two cases.

## 3. Results & Discussion

### 3.1 Testing of existing silver(I) parameters on HEFM tetrapeptide

The silver(I) binding constants obtained for the HEFM tetrapeptide prior the calibration procedure are reported in Table 1. The values are given for different cases, combining box size (30 and 50 Å, to verify potential size effects) and parameters sets (Won, *HFE* and *IOD*). In all cases we can notice that the binding constants are lower or equal to 10 (log_10_(*K*_*bind,calc*_) ≤ 1), indicating a drastic underestimation of the interaction strength between silver(I) ion and the HEFM peptide (experiment: log_10_(*K*_*bind,exp*_)=6.6).^8^ This is not really surprising because those parameters were developed to model silver(I)-water interaction only, without any intended correction for interactions with organic solutes.

### 3.2 Testing of existing parameters on sidechain fragments and calibration of NBFIX parameters

When the *HFE* parameters of Merz^12^ were tested on the interaction of silver(I) with the sidechain fragments (Table 2, column “*HFE*”) we note that the interaction strength is underestimated for all sidechain fragments except for propanoate. For the latter the calculated binding constants were reasonably close to the experimental value (0.89 and 1.25 versus 0.73, respectively for cTIP3P and sTIP3P); the same holds for the equilibrium distance (see also ESI for the PMF profile).

The NBFIX parameters optimized to reproduce the experimental binding constant of the sidechains fragments are reported in Table 2, together with the calculated and experimental binding constants. The fragments 4-ethylimidazole and ethylmethylsulfide possess the highest experimental binding constants of 3.85 and 3.6, respectively (in logarithmic scale). This can explain the high binding constants of the peptides containing such residues (log_10_(*K*_*bind,exp*_)=5-6.7). In the CHARMM36m force field the sulfur of methionine is basically charge neutral (−0.09 *e*) which means that there is no electrostatic stabilization for the interaction with silver(I). Thus, a very strong NBFIX correction (high *ε*^*NBFIX*^ in absolute terms) needs to be applied on the sulfur atom of ethylmethylsulfide in order to correct its affinity. This result can be better appreciated observing Figure 3, which contains the PMF profiles of three sidechain fragments before and after the calibration (in black and red lines, respectively). As a general observation, the *HFE* parameter set yields less attractive PMFs. In particular, in the case of ethylmethylsulfide and toluene, the interaction passes from purely repulsive to attractive after the calibration procedure, which explains why the binding constant of the HEFM peptide was drastically underestimated during the preliminary evaluations.

**Figure 3.**
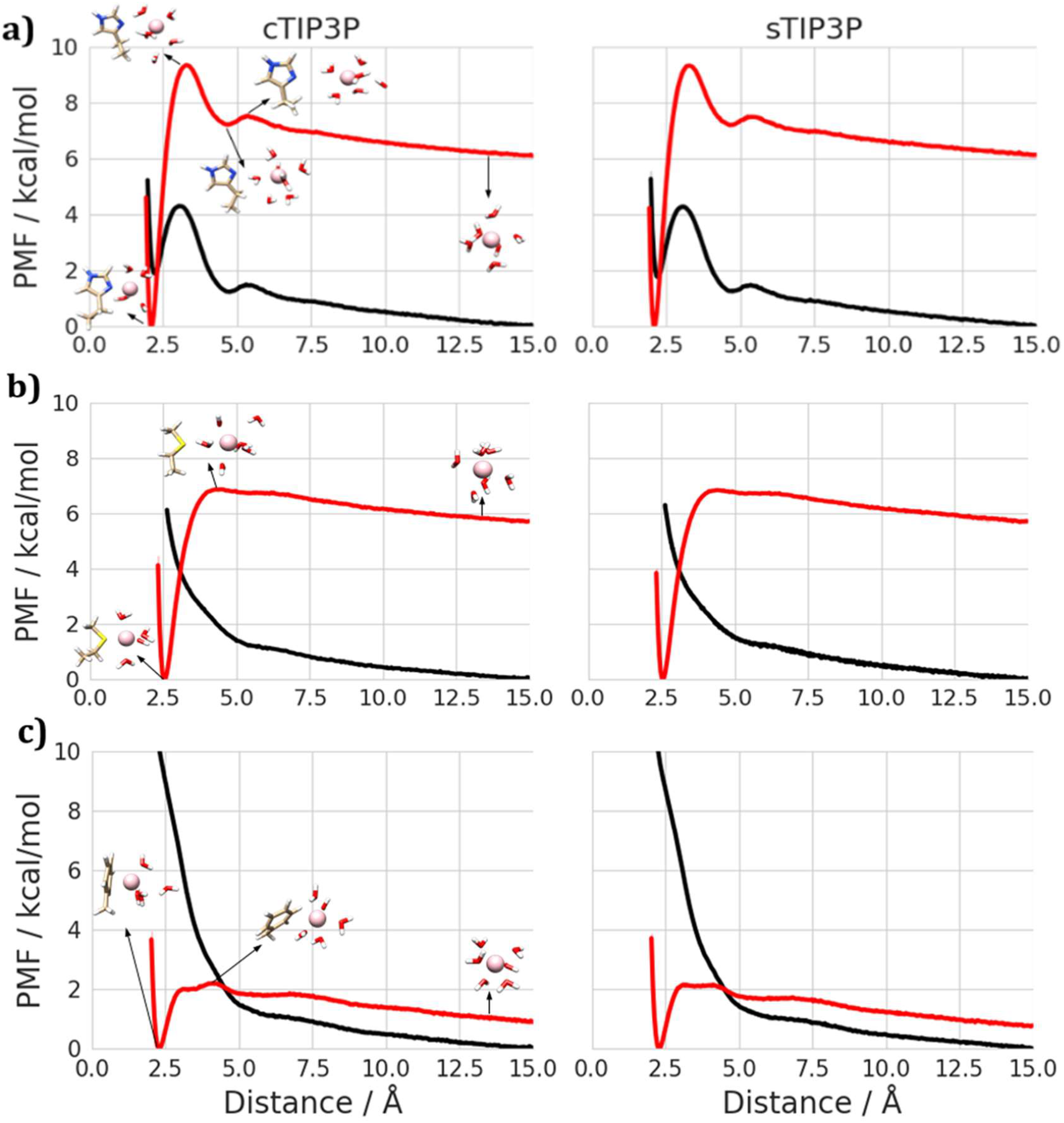
PMF profiles and structures of (a) 4-ethylimidazole, (b) ethylmethylsulfide and (c) toluene with *HFE* only parameters (black line) and with NBFIX parameters (red line) and in cTIP3P (left) and sTIP3P (right) water models. PMF curves are plotted together with the standard error of the mean obtained from an average over four blocks (lighter colors, barely visible due to the small magnitude). For propanoate, the distance refers to the one from the carboxylic C atom. For toluene, the distance refers to the one from the center of the aromatic ring. Structures are reported for the cTIP3P water model.

### 3.3. Validation on four silver-binding tetrapeptides

To validate the fragment-optimized NBFIX parameters, we calculated the binding constants for the silver-binding tetrapeptides HEFM, MNEH, HAAM and MAAH (Table 3). Overall, the calculated binding constants are on the same order of magnitude as the experimental values. In terms of Gibb’s free energy, they are within 1 kcal/mol of the experimental reference values. Moreover, no significant difference is found when comparing cTIP3P and sTIP3P.

**Table 3.**
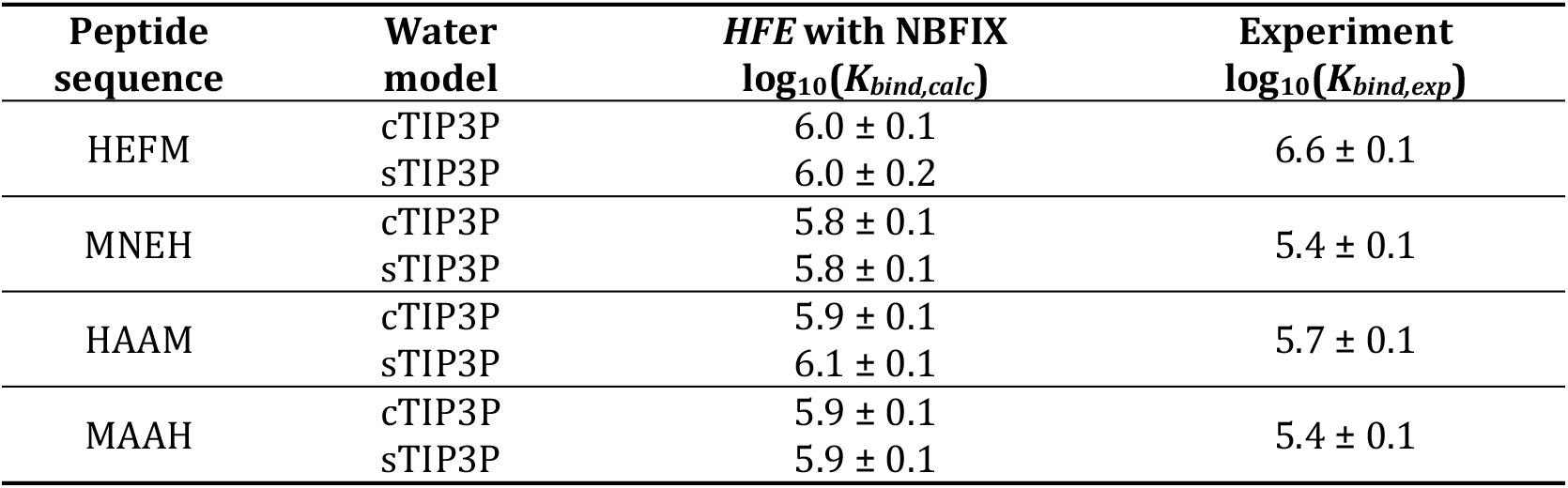
Calculated binding constants for the studied tetrapeptides (HEFM, MNEH, HAAM and MAAH) with the optimized NBFIX paramters. For each peptide eight potential-scaled MD simulations of about 800 ns each were performed (see ESI for more details). The results are shown for both cTIP3P and sTIP3P. Calculated association constants are reported together with the standard error of the mean from block analysis.

| Peptide sequence | Water model | HFE with NBFIX<br>$\log_{10}(K_{bind,calc})$ | Experiment<br>$\log_{10}(K_{bind,exp})$ |
| --- | --- | --- | --- |
| HEFM | cTIP3P | $6.0 \pm 0.1$ | $6.6 \pm 0.1$ |
| | sTIP3P | $6.0 \pm 0.2$ | |
| MNEH | cTIP3P | $5.8 \pm 0.1$ | $5.4 \pm 0.1$ |
| | sTIP3P | $5.8 \pm 0.1$ | |
| HAAM | cTIP3P | $5.9 \pm 0.1$ | $5.7 \pm 0.1$ |
| | sTIP3P | $6.1 \pm 0.1$ | |
| MAAH | cTIP3P | $5.9 \pm 0.1$ | $5.4 \pm 0.1$ |
| | sTIP3P | $5.9 \pm 0.1$ | |

Whereas the experiments measure subtle differences between these peptides in terms of affinity, the calculations predict basically the same binding constant for all peptides. A potential reason for this levelling/flattening is the neglecting of the protonation/tautomerism of histidine in the calculations above. Here we would like to remind the reader that calculated binding constants of Table 2 correspond basically to microscopic binding constants for specific peptides where histidine can only exist in its tautomer Hse; Hse is the preferred silver(I)-binding tautomer for small peptides (see above). According to constant-pH simulations of the apo form, the MAAH peptide displays a significantly decreased population of the preferred Hse tautomer (36%) with respect to HAAM (60%). Thus, there is a larger tautomeric penalty to overcome for MAAH than for HAAM; see the companion article for more details on this tautomeric correction.^7^ As a result, the calculated binding constant with tautomeric correction, log_10_(*K*_*bind,corr*_), is 0.22 lower for MAAH (5.47) than for HAAM (5.69) in excellent agreement with the experiment (5.4 ± 0.1 and 5.7 ± 0.1 for MAAH and HAAM, respectively).

### 3.4 Application to oligopeptide LP1

As a further validation of the parameters, we tested the ability to reproduce structural features of silver-bound peptides, i.e., the increase of *α*-helical content of the 14-residue peptide LP1 in the presence of silver(I) ions.^9^ Figure 4 reports the average α-helical content of the LP1 peptide (in percentage) as a function of simulation time; at the beginning of the simulation the peptide was completely extended (linear backbone). A single silver(I) ion in the simulation box is sufficient to increase the *α*-helical content to about 17-18% within 5 μs in the case of Jorgensen’s water model (sTIP3P; Figure 4b). By contrast, in absence of the ion the *α*-helical content remains around 5-6%. With the CHARMM TIP3P water model no such increase is seen for a single silver(I) ion. When the number of silver(I) ions is, however, augmented to 5 the helical content increases, too, with respect to the simulation without silver(I) ions in agreement with the experiment. It seems that the sTIP3P water model induces more easily α-helical motifs in agreement with previous observations.^34^

**Figure 4.**
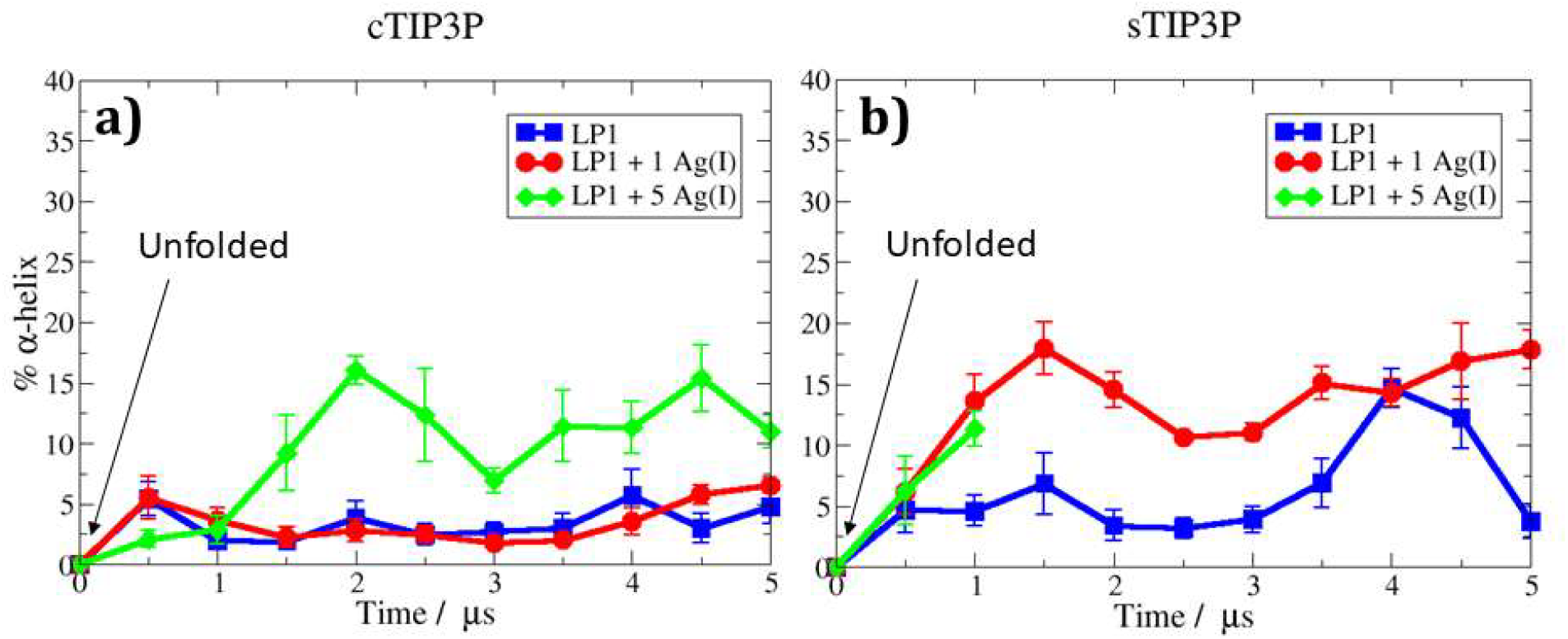
Percentage of *α*-helical content of LP1 over time (µs) for simulations in cTIP3P (a) and in sTIP3P (b) in the absence (apo, blue) and presence of 1 (red) or 5 (green) silver(I) ions.

Overall the α-helical content is still relatively modest after 5 μs of MD. It should be noted that folding of the peptide into a helix is a slow process and our simulation time could not capture the full dynamics. A part of the problem is that the chosen simulation approach (with a scaled potential) slows down the folding dynamics since the scaling favors the silver(I) dissociation.

## 4. Conclusion & Outlook

Our new force field parameters for silver(I) ions are capable to reproduce binding constants for tetrapeptides on the same order of magnitude as experimental values. Also, the increased α-helical content of LP1 in the presence of silver(I) was correctly detected with these new parameters. These results validate the applicability of the simple yet accurate LJ model, which can be tightly tuned by means of the NBFIX feature of the CHARMM force field family to selectively correct atom-atom non-bonded interactions, without affecting the rest of the force field. We anticipate that the methodology proposed in this work can be in principle applied to any metal ion provided the existence of reference data for binding constants and distances and the validity of the LJ potential to represent that ion.

We note, however, further improvements for the future simulations of silver(I) interactions with peptides and proteins:

- Histidine tautomerism. The explicit inclusion of the various protonation states of histidine (and in general on all titrable amino acids, especially for cysteine). In this work we assumed that a single static protonation state (Hse) with a microscopic binding constant that coincides with the experimental macroscopic binding constant of 4-ethylimidazole. A more realistic description would allow a dynamic change of histidine protonation states by constant-pH simulations^35–37^ where each state binds silver(I) with a specific microscopic binding constant.
- Polarization. This effect is particularly pertinent to deal with the coordination of three or more amino acids to the same silver(I) center as it is observed in more complex systems such as silver-binding oligopeptides (LP1) or proteins (SilE). The NBFIX LJ potentials are fully additive and therefore may lead to an overestimation of the affinity. Inclusion of polarization could be done in the context of a polarizable force field such as AMOEBA or CHARMM’s Drude force field.^38,39^ Another potential limitation of 12-6 potential due to the lack of polarizability could affect the conformation transitions involving polar secondary structure elements (e.g. α-helices).
- Cationic Dummy Metal Ion Model. To account for particular coordination geometries, the extension to a model with cationic dummy charges could be beneficial.^40^
- Extension of the parametrization to other sidechain fragments (e.g., thiols/thiolates to represent cysteine) and the peptide backbone. The availability of experimental data is the limited factor here. In this light, we would like to emphasize that a careful determination of the reference parameters (equilibrium distance and binding constant) is crucial for the parametrization procedure and can highly influence the sampling.
- Sampling. Finally, improvements may result through the use of enhanced sampling techniques such as Replica Exchange.^41^ This could be crucial for an integrative approach that combines the affinity simulation of silver(I) with the constant-pH feature.

### Electronic Supporting Information

ESI is available online at https://doi.org/10.1101/2024.11.28.625676. It contains experimental methods and computational details.

## Supporting information

Electronic Supplementary Information

## Acknowledgment

The authors would like to acknowledge the High Performance Computing Center of the University of Strasbourg for supporting this work by providing scientific support and access to computing resources through grants g2023a142c/g and g2024a236c/g. Part of the computing resources were funded by the Equipex Equip@Meso project (Programme Investissements d’Avenir) and the CPER Alsacalcul/Big Data. The work was further supported by a grant from the Swiss National Science Foundation (Project N° 204215). LM is grateful to the Université de Haute Alsace for a PhD stipend. A.B. and K.F. thank the University of Fribourg and the

## List of abbreviations & symbols

PB1b: oligopeptide Ac-AHQKMVESHQRMMG-NH_2_
CA: CHARMM36m atom type for aromatic carbon of amino acids
CD: Circular Dichroism
CHARMM: Chemistry at HARvard Macromolecular Mechanics
cTIP3P: CHARMM’s TIP3P water model
*D*: product of the vacuum permittivity and the relative permittivity
Δ*G*_*bind,calc*_: Calculated binding free energy
DFT: Density Functional Theory
*ε*: well-depth of the Lennard-Jones potential
*E*_*elec*_ *& E*_LJ_: electrostatic & Lennard-Jones interaction energies
*E*_*NB* J_: non-bonded interaction energy
*d*_*eq,calc*_ *& d*_*eq,target*_: calculated and target equilibrium distance
*γ*_*d*_, *γ*_*K*_: pairwise correction coefficients Glu: glutamic acid
HAAM: tetrapeptide Ac-HisAlaAlaMet-NH_2_
HEFM: tetrapeptide Ac-HisGluPheMet-NH_2_
*HFE*: hydration free energy
His: histidine
Hsd: δN-tautomer of histidine
Hse: *ε*N-tautomer of histidine
Hsp: protonated histidine
*IOD*: ion-oxygen distance
*k*_*B*_: Boltzmann constant
*K*_*bind,calc*_ *& K*_*bind,exp*_: calculated and experimental binding constant
*K*_*bind,corr*_: calculated binding constant with tautomeric correction
*k*_*ij*_: binary interaction coefficient for *ε*
*l*_*ij*_: binary interaction coefficient for *R*^*min*^
LJ: Lennard-Jones
LP1: see B1b
MAAH: tetrapeptide Ac-HisAlaAlaMet-NH_2_
MD: molecular dynamics
Met: methionine
MNEH: tetrapeptide Ac-MetAsnGluHis-NH_2_
NB: non-bonded
NBFIX: non-bonded fix
NMR: Nuclear Magnetic Resonance
NR2: CHARMM36m atom type for non-protonated imidazole nitrogen of histidine
Phe: phenylalanine
PMF: potential of mean force
*Q*: product of partial charges of two interaction atoms
*r*: interatomic distance
*R*^*min*^: distance where the Lennard-Jones potential takes a minimum
S: CHARMM36m atom type for methionine sulfur
sTIP3P: Jorgensen’s TIP3P water model
*T*: Temperature
TIP3P: transferable intermolecular potential with 3 points
US: Umbrella sampling
WHAM: weighted histogram analysis method

