## Supplementary material for "Modelling the interaction between silver(I) ion and proteins with the 12-6 Lennard-Jones potential: a bottom-up parameterization approach": Electronic Supplementary Information

#### Index

### 1. Experimental determination of the binding constants of imidazole, 4(5)-methylimidazole 4(5)-ethylimidazole and ethylmethylsulfide

#### 1.1 Starting materials, solvents and instrumentation

*Potentiometry:* High purified water (Honeywell Chromasolv™ plus) was de-oxygenated by $CO_2$- and $O_2$-free argon just before use. The stock solutions of **imidazole** (Merck), **4(5)-methylimidazole** (Sigma Aldrich, purum p.a., 98%) and **4(5)-ethylimidazole** (BLD Pharmatech GmbH, purum p.a. 98%) were prepared by weighing solids using an METTLER TOLEDO XA105 analytical balance (accuracy 0. 1 mg). The ionic strength was maintained at 0.1 M with sodium perchlorate ($NaClO_4$, Merck kGaA), and all measurements were carried out at 25.0°C. The silver(I) stock solutions (~ 5 x $10^{-2}$ M) were prepared by dissolving appropriate amounts of solid Ag(I) perchlorate ($Ag(ClO_4)$, Sigma Aldrich, purum p.a., 97%) in water and store in dark.

*Spectrophotometry*: $AgClO_4$ and **ethylmethylsulfide** (Thermo Scientific Chemicals, 97%) were dissolved with boiled and $CO_2$-free 18 MΩ.

#### 1.2 Potentiometric titrations

The potentiometric titrations of the free ligands **imidazole** (3,88.$10^{-3}$ M) and **4(5)-ethylimidazole** (4,72.$10^{-3}$ M) and their silver complexes (4.5 ≤ $[\text{ligand}]_{tot}/[Ag^+]$ < 10) were performed using an automatic titrator system 794 Basic Titrino (METROHM) with a combined glass electrode (METROHM 6.0234.500, Long Life) filled with 0.1 M NaCl in water and connected to a microcomputer (TIAMO light 1.2 program). The combined glass electrode was calibrated as a hydrogen concentration probe by titrating known amounts of perchloric acid (0,096 M from $HClO_4$, MERCK, puriss p.a. 99,9%, >70 %) with $CO_2$-free sodium hydroxide solution (~ 0,109 M from NaOH, Alfa Aesar, 97%). The $HClO_4$ and NaOH solutions were freshly prepared just before use and titrated with sodium tetraborate decahydrate ($B_4Na_2O_7.10H_2O$, FLUKA, puriss, p.a., > 99.5%) and potassium hydrogen phthalate ($C_8H_5KO_3$, SIGMA ALDRICH, p.a., > 99.5%), respectively, with methyl orange (RAL) and phenolphthalein (PROLABO, purum) used as colorimetric indicators. The temperature of the titration cell was maintained at 25.0 ± 0.2 °C with the help of a LAUDA E200 thermostat. The GLEE program[1] was applied for the glass electrode calibration (standard electrode potential $E_0$/mV and slope of the electrode/mV $pH^{-1}$) and to check carbonate levels of the NaOH solutions used (< 5 %). The potentiometric data of ligands and their silver complexes (about 300 points collected over the pH range 2.5-11.5) were refined with the HYPERQUAD 2000[2] program which uses non-linear least-squares methods.[3] Potentiometric data points were weighted by a formula allowing greater pH errors in the region of an end-point than elsewhere. The weighting factor $W_i$ is defined as the reciprocal of the estimated variance of measurements: $W_i = 1/\sigma_i^2 = 1/[\sigma_E^2 + (\delta E/\delta V)^2\sigma_V^2]$ where $\sigma_E^2$ and $\sigma_V^2$ are the estimated variances of the potential and volume readings, respectively. The constants were refined by minimizing the error-square sum, *U*, of the potentials:

$$U = \sum_{i}^{N} W_i (E_{obs,i} - E_{cal,i})^2$$

At least three or four potentiometric titrations were treated as single sets, for each system. The quality of fit was judged by the values of the sample standard deviation, *S*, and the goodness of fit, $\chi^2$, (Pearson's test). At $\sigma_E$ = 0.1 mV (0.023 $\sigma_{pH}$) and $\sigma_V$ = 0.005 mL, the values of *S* in different sets of titrations were between 0.6 and 1.6, and $\chi^2$ was below 99. The scatter of residuals *vs.* pH was reasonably random, without any significant systematic trends, thus indicating a good fit of the experimental data. The stability and successive protonation constants were calculated from the cumulative constants determined with the program. The uncertainties in the $\log_{10}(K)$ values correspond to the added standard deviations in the cumulative constants.

#### 1.3 Acid-base properties of ligands

Protonation constant (charges are omitted):

$L + H \rightleftarrows LH$ ;  $\log_{10}(K_{11})$

**Table S1:** Logarithm of the protonation constant for ligands **imidazole** and **4(5)-ethylimidazole** compared to literature data reported for closely related systems.

| Ligand | imidazole, $L_1$ | 4(5)-methylimidazole, $L_2$ | 4(5)-ethylimidazole, $L_3$ |
|---|---|---|---|
| $\log_{10}(K_{11})$ | 7.09 ± 0.03 | 7.64 ± 0.02 | 7.59 ± 0.01 |
| F. Peral, 1997[4] | 6.99 | 7.56 | / |
| S. Nakatsuji, 1969[5] | 7.33 (1M $KNO_3$) | / | / |

For ligands $L_1$, $L_2$ and $L_3$ each solution had a pH of 8.5. Then, for each measurement, an acid solution was added to start the measurement at acidic pH.

#### 1.4 Potentiometry of ligand imidazole

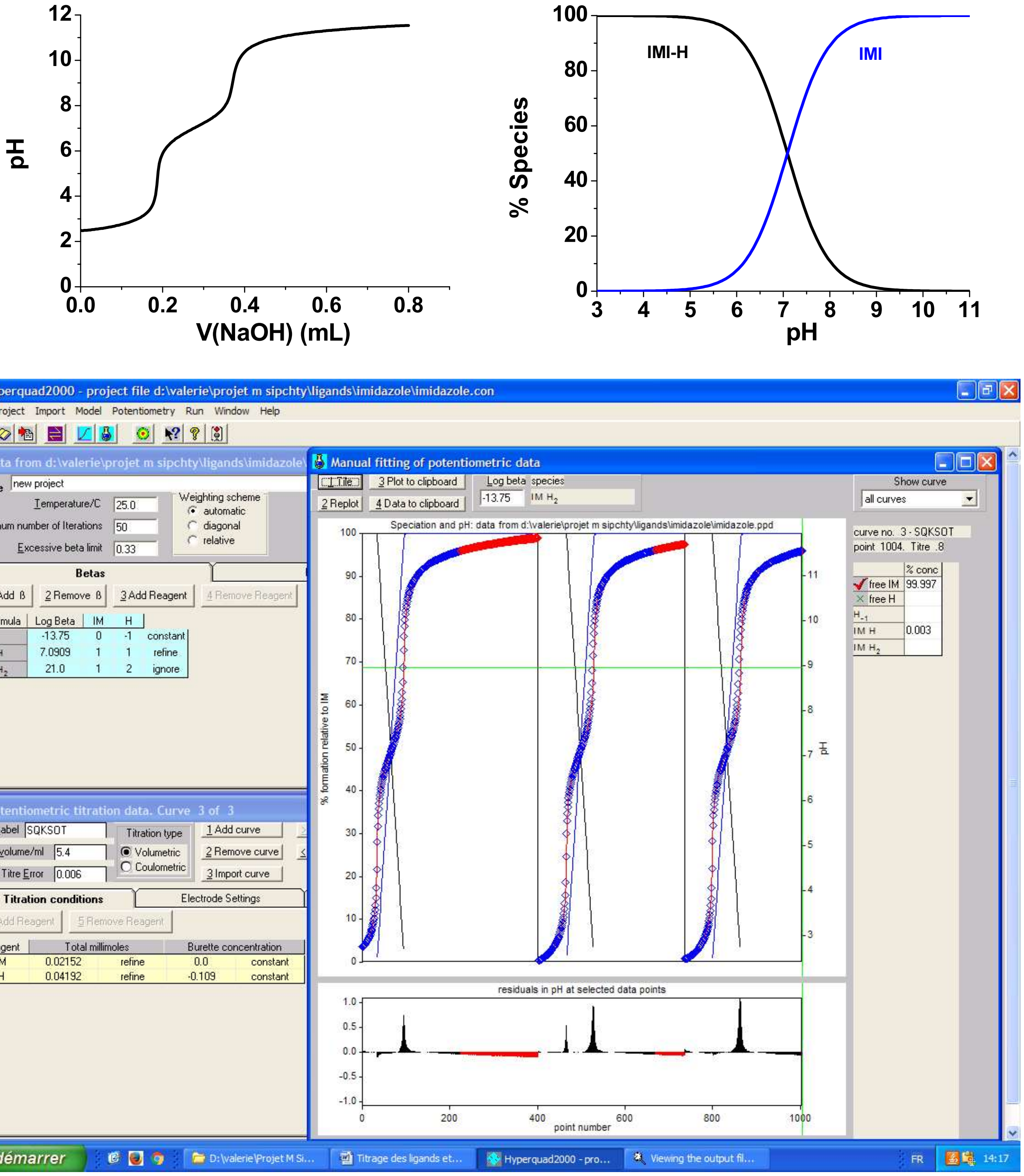

**Figure S1.** Top: Potentiometric titration curve of **imidazole** (left) and distribution diagram (right; from hyss simulation). Bottom: Hyperquad analysis. Solvent: Water; I = 0.1 M $NaClO_4$; T = 25°C; [**imidazole**]0 = 3.88 x $10^{-3}$ M. σ = 1.17.

#### 1.5 Potentiometry of ligand 4(5)-methylimidazole

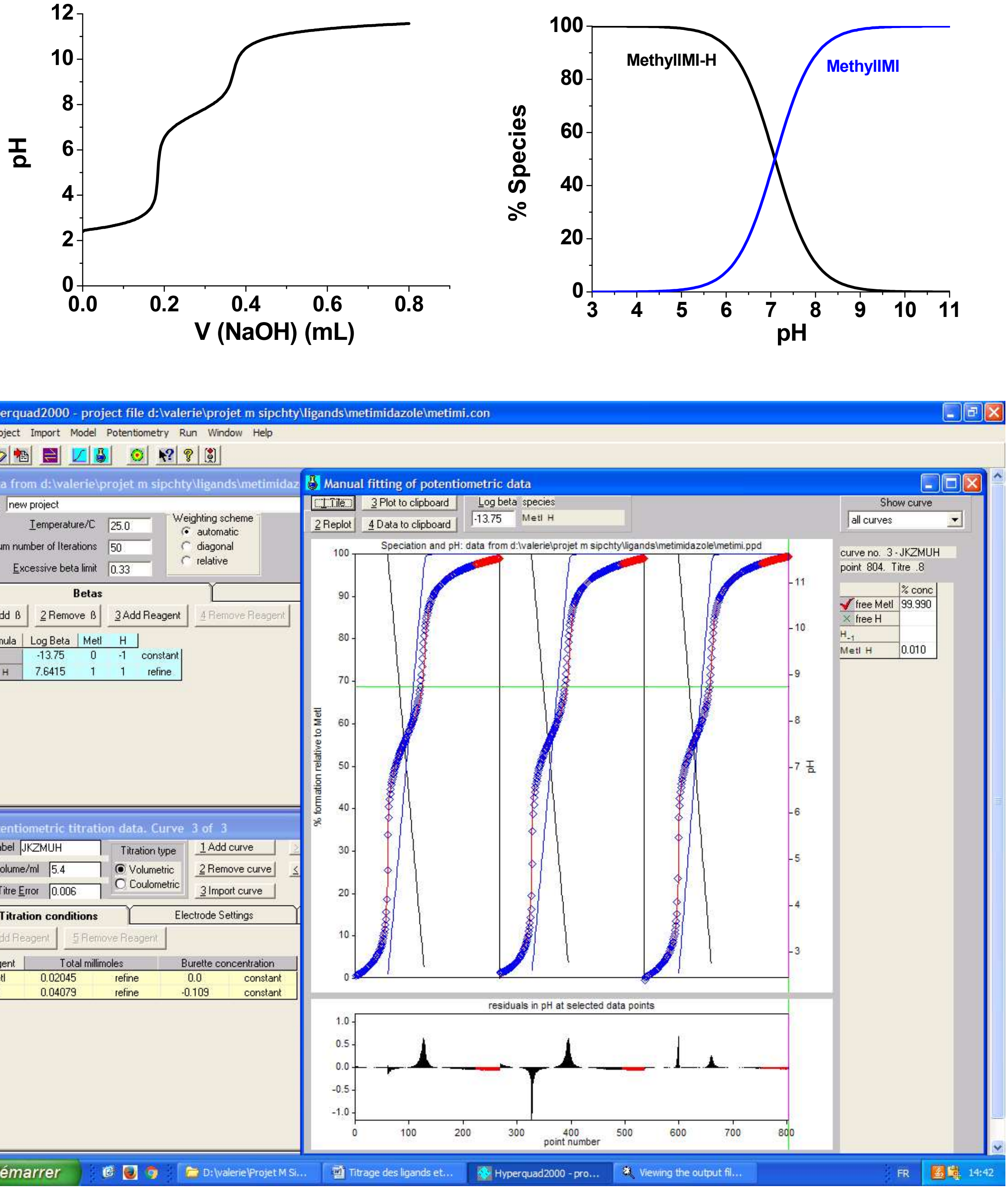

**Figure S2.** Top: Potentiometric titration curve of **4(5)-methylimidazole** (left) and distribution diagram (right; from hyss simulation). Bottom: Hyperquad analysis. Solvent: Water; I = 0.1 M $NaClO_4$; T = 25°C; [**4(5)-methylimidazole**]0 = 4.07 x $10^{-3}$ M. σ = 1.06 (selected range in blue).

#### 1.6 Potentiometry of ligand 4(5)-ethylimidazole

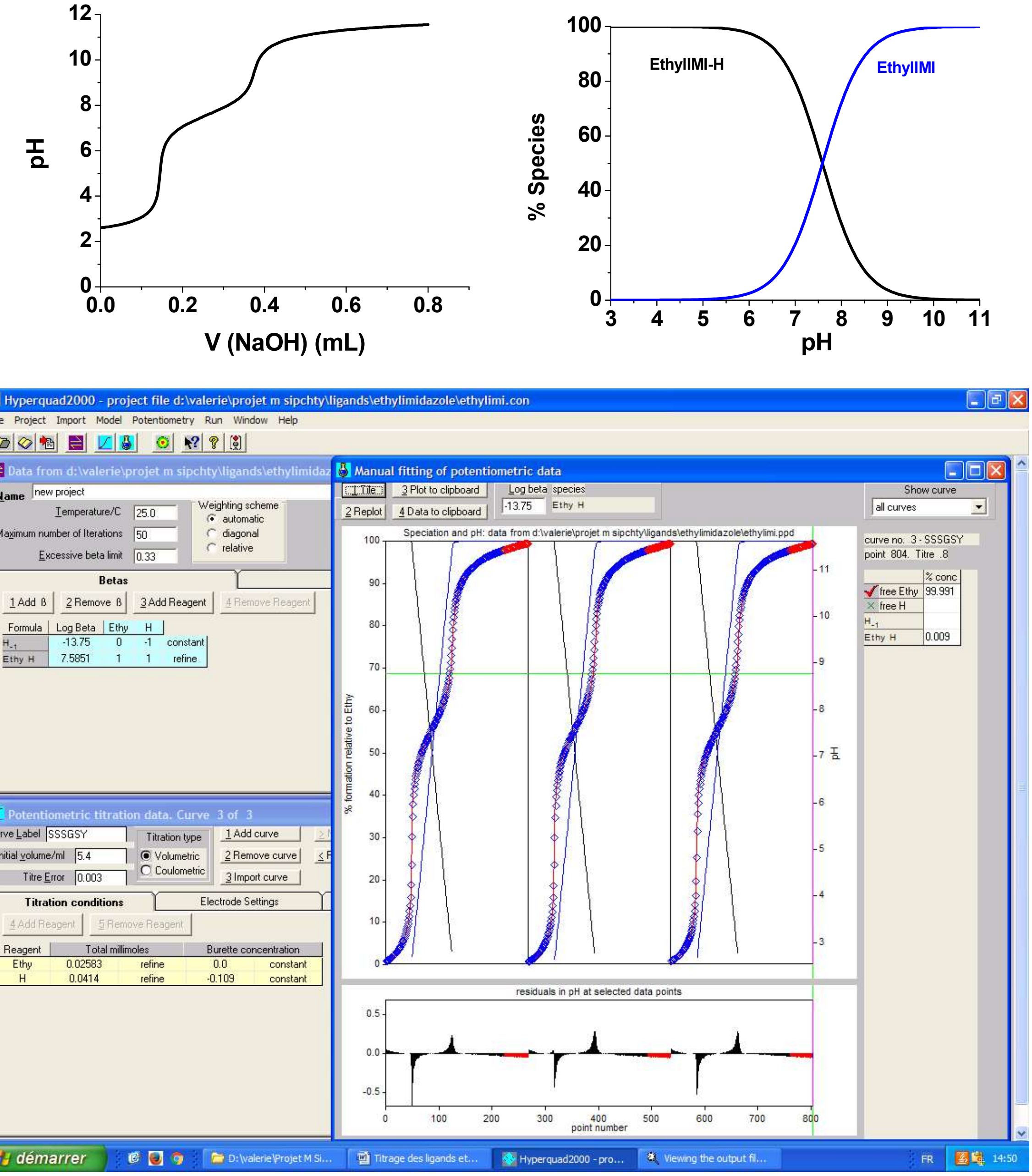

**Figure S3.** Top: Potentiometric titration curve of **4(5)-ethylimidazole** (left) and distribution diagram (right; from hyss simulation). Bottom: Hyperquad analysis. Solvent: Water; I = 0.1 M $NaClO_4$; T = 25°C; [**4(5)-ethylimidazole**]0 = 4.72 x $10^{-3}$ M. σ = 1.02.

#### 1.7 Determination of the binding constants with silver(I)

The studied additional equilibrium reactions were:

Ag + L ⇄ AgL   $\log_{10}(\beta_{11})$

Ag + 2L ⇄ $AgL_2$   $\log_{10}(\beta_{12})$

**Table S2:** Logarithms of the stability constants for the silver complexes with ligands **imidazole, 4(5)-ethylimidazole** and **ethylmethylsulfide** compared to literature data reported for closely related systems.

| Ligand | imidazole, $L_1$ | 4(5)-methylimidazole, $L_2$ | 4(5)-ethylimidazole, $L_3$ | ethylmethylsulfide, $L_4$ |
|---|---|---|---|---|
| $\log_{10}(\beta_{11})$ | 3.2 ± 0.1 | 3.75 ± 0.08 | 3.85 ± 0.04 | 3.6 ± 0.3 |
| $\log_{10}(\beta_{12})$ | 7.08 ± 0.06 | 7.4 ± 0.1 | 7.23 ± 0.09 | 6.1 ± 0.4 |
| S. Nakatsuji, 1969[5] | $\log_{10}(\beta_{11})$ = 3.08<br>$\log_{10}(\beta_{12})$= 6.95 | | | |
| D. H. Gold, 1960[6] | $\log_{10}(\beta_{11})$ = 3.11<br>$\log_{10}(\beta_{12})$ = 6.84 | | | |
| J. E. Bauman, 1963[7] | $\log_{10}(\beta_{11})$ = 3.05<br>$\log_{10}(\beta_{12})$ = 6.88 | | | |

Each initial solution had a pH=8.5. For each measurement, acid was added following Ag ions to have an acid pH at the start of the titration. Some colloidal species were formed at neutral pH. Possibly it was $[Ag_3(Ligand)_6][ClO_4]_3$.[8] The precipitated constants of $Ag^+$ in basic media $\log_{10}(K_{Ag2O})$= -17.7 have been considered throughout the processing of the potentiometric data.

##### 1.7.1 Ag+ coordination properties of ligand imidazole

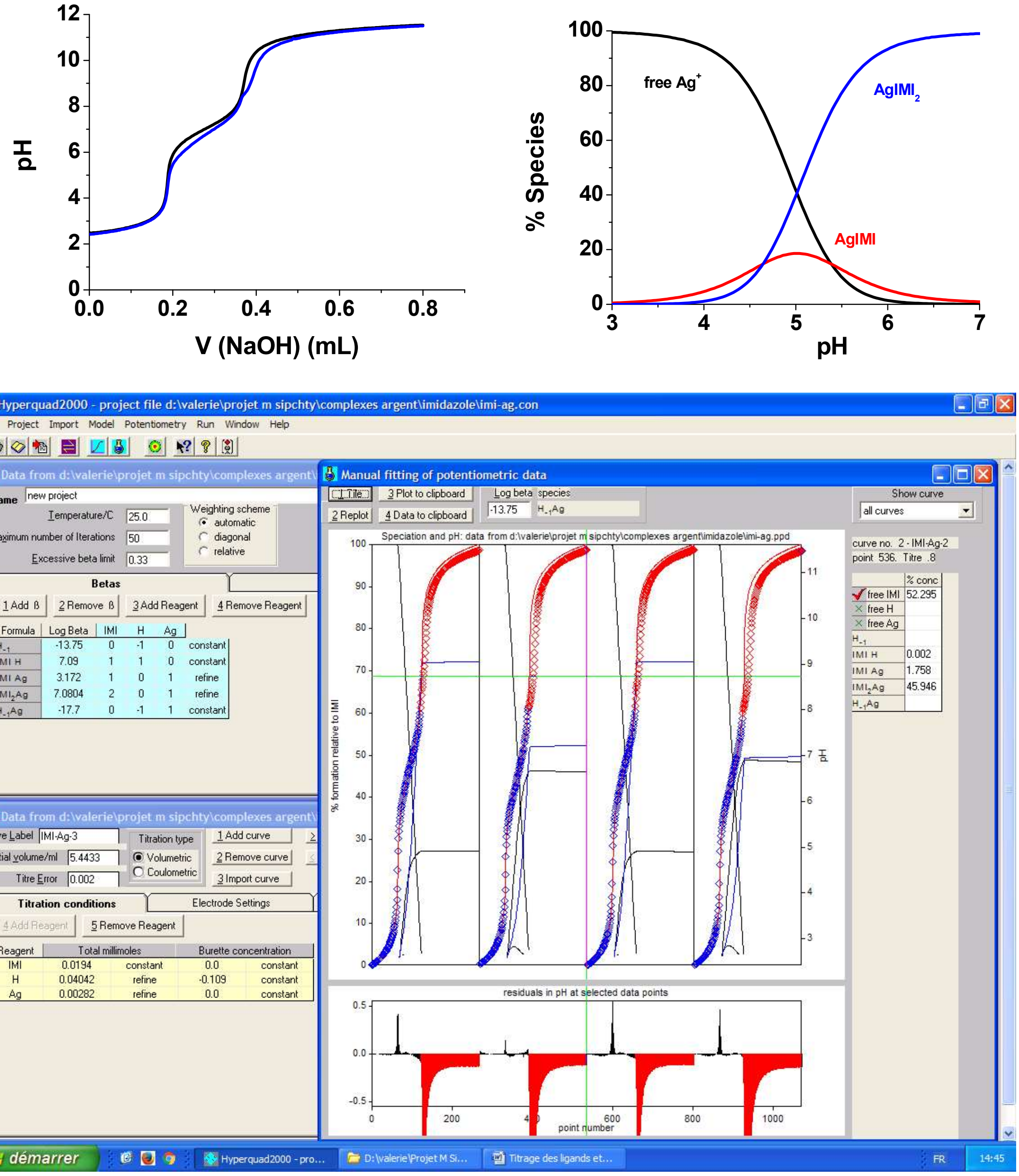

**Figure S4.** P Top: Potentiometric titration curve of **imidazole** (top black curve, left) and **Ag/imidazole** (top blue curve, left) and distribution diagram (right). Bottom: Hyperquad analysis (bottom). Solvent: Water; *I* = 0.1 M $NaClO_4$; *T* = 25°C; [**imidazole**]$_0$ = 3.88 x $10^{-3}$ M; [Ag]$_0$ =4 .02.$10^{-4}$ or 8.00.$10^{-4}$ M. σ = 0.64 (selected range in blue).

#### 1.7.2 $Ag^+$ coordination properties of ligand 4(5)-methylimidazole

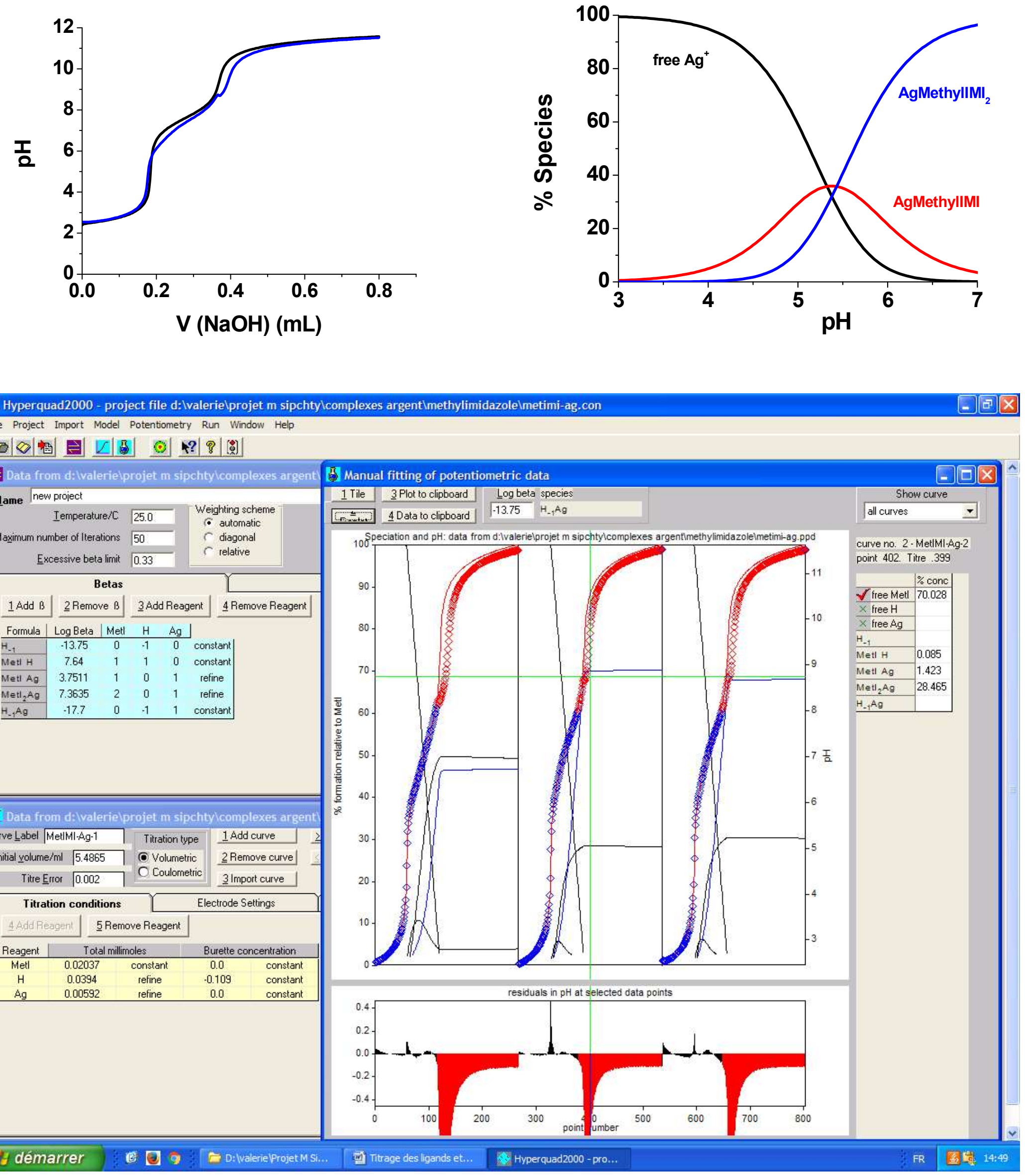

**Figure S5.** Top: Potentiometric titration curve of **4(5)-methylimidazole** (top black curve, left and Ag/**4(5)-methylimidazole** (top blue curve, left) and distribution diagram (right; from hyss simulation). Bottom: Hyperquad analysis (bottom). Solvent: Water; $I$ = 0.1 M $NaClO_4$; $T$ = 25°C; $[\textbf{4(5)-methylimidazole}]_0 = 4.07 \times 10^{-3}$ M; $[Ag]_0$ =4 .02.$10^{-4}$ or 8.00.$10^{-4}$ M. σ = 0.79 (selected range in blue).

##### 1.7.3 $Ag^{+}$ coordination properties of ligand 4(5)-ethylimidazole

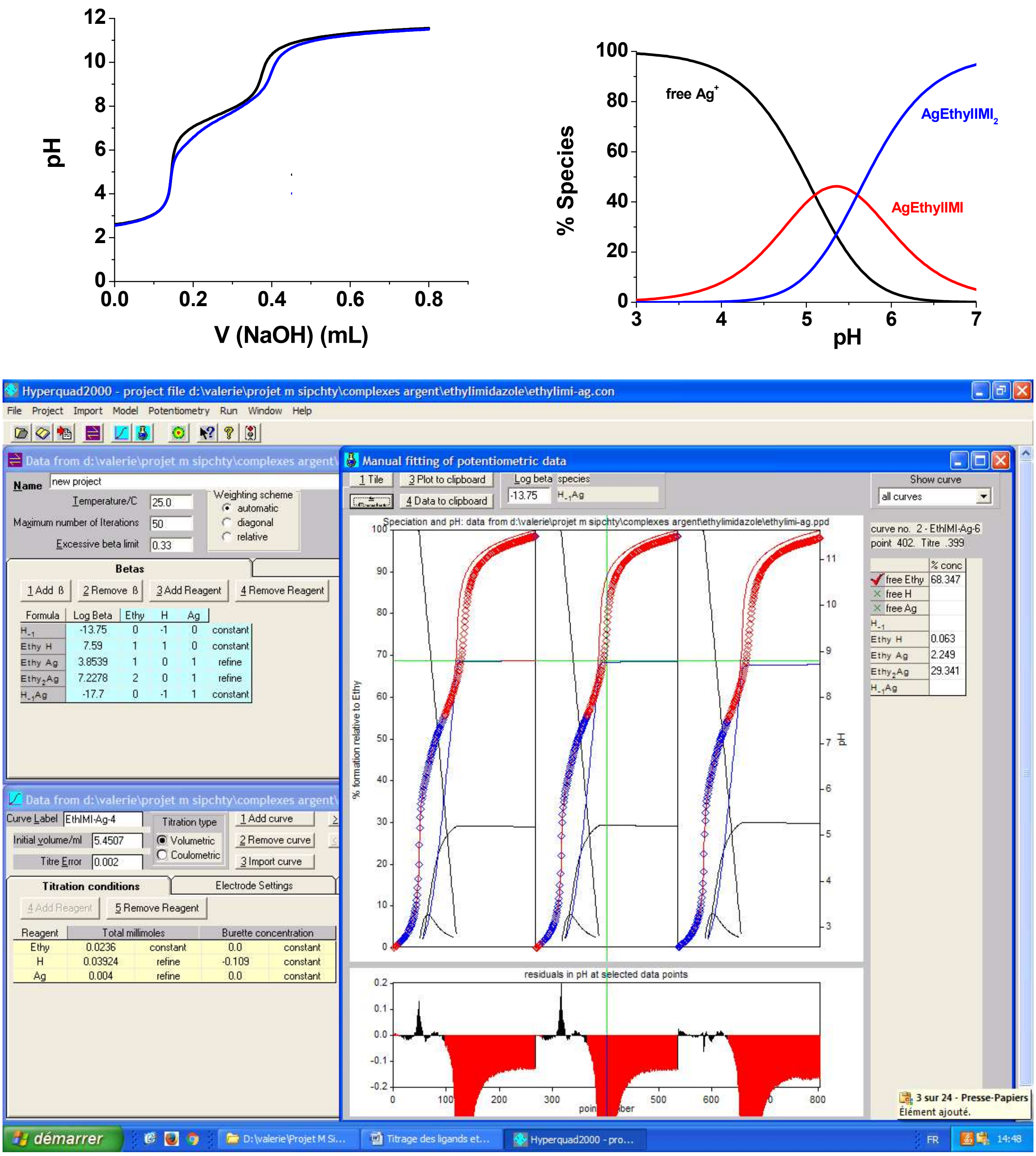

**Figure S6.** Top: Potentiometric titration curve of **4(5)-ethylimidazole** (top black curve, left) and **Ag/4(5)-ethylimidazole** (top blue curve, left) and distribution diagram (right; from hyss simulation). Bottom: Hyperquad analysis (bottom). Solvent: Water; $I$ = 0.1 M $NaClO_4$; $T$ = 25°C; [**4(5)-ethylimidazole**]$_0$ = 4.72 x $10^{-3}$ M; $[Ag]_0$ =4 .70.$10^{-4}$ M. σ = 0.47 (selected range in blue).

##### 1.7.4 $Ag^{+}$ coordination properties of ligand ethylmethylsulfide

*Spectrophotometric properties*: Since protonation on **ethylmethylsulfide** does not occur the direct potentiometric titration with NaOH cannot be used. Instead we investigated the spectrophotometric of silver(I) and **ethylmethylsulfide**. Silver(I) ion possesses two absorption bands at 210 and 225 nm and **ethylmethylsulfide** one at 207 nm. The complex shows a band at 231 nm.

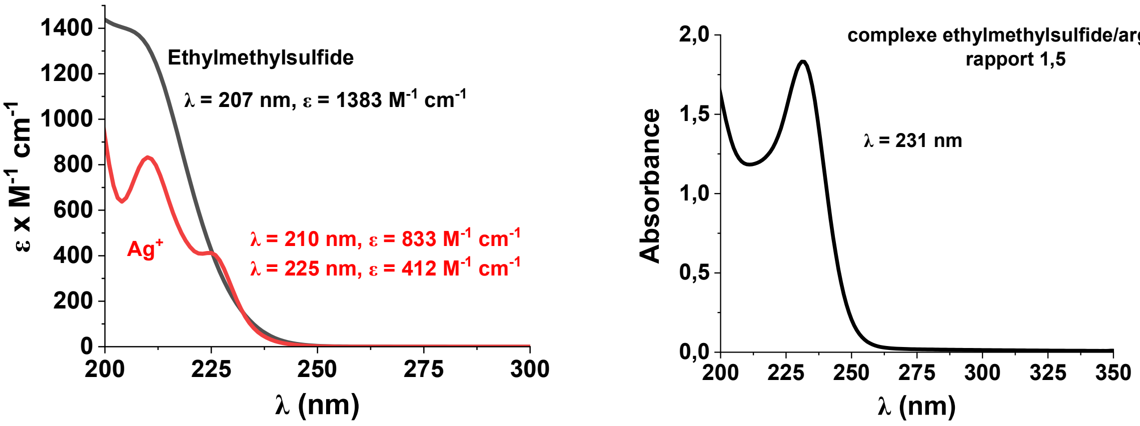

**Figure S7.** Electronic absorption spectra of $Ag^+$, **ethylmethylsulfide** and absorption spectrum of silver complexes with ligand $L_4$ at pH = 5.50.

*Silver ion titrations with ethylmethylsulfide:* The titrations were performed by sequencial additions of **ethylmethylsulfide** into the Ag solution. Thereby the sulfide was taken directly from the volumetric flask and added to the silver solution, then the absorbance spectrum is immediately measured. All the measurement were carried on spectrophotometer Agilent Cary 5000 with quartz cuvette with 0.2 cm optical path length. The UV absorption titrations of Ag+ ions and **ethylmethylsulfide** were performed at pH=3. The absorption spectrophotometric titration *versus* **ethylmethylsulfide** concentration has been processed with the Specfit program.[3,9–13]

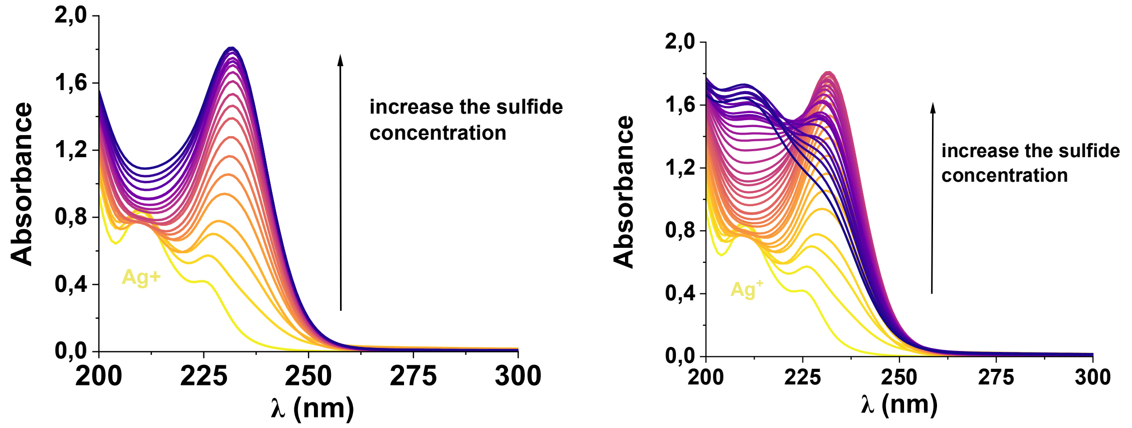

**Figure S9.** UV absorption spectra as a function of $L_4$ concentration with silver ion at pH 3.50, solvant : $H_2O$; $T$ = 25.0°C; $[Ag^+]_{init}$ = $5.10^{-3}$ M, $[L_4]_{init}$ = 7.5 x $10^{-3}$ M. The absorption spectra have not been corrected from dilution effects.

The analysis of the first 20 spectra with Specfit, leaving the absorption spectrum of silver(I) and **ethylmethylsulfide** as "unknown", gives $\log_{10}(\beta_{11})$ = 3.6 ± 0.3.

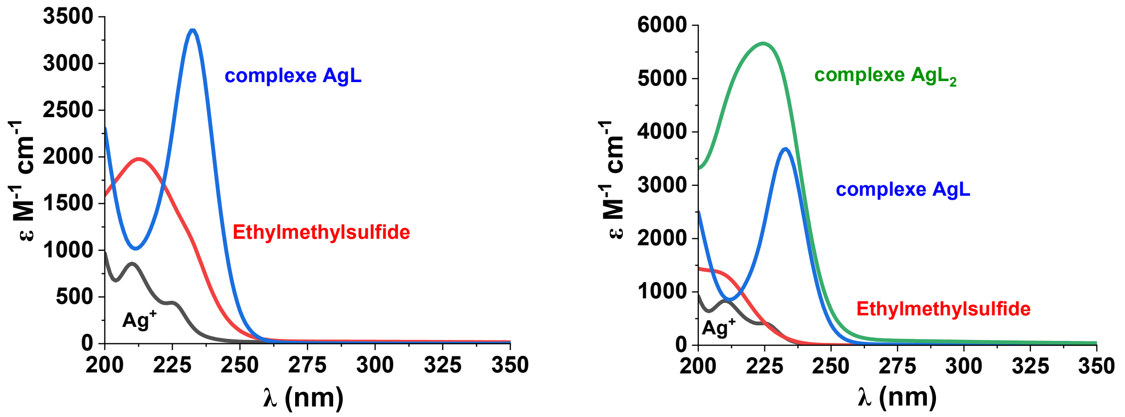

**Figure S10.** Apparent electronic absorption spectra of $Ag^{+}$, **ethylmethylsulfide** and its silver complexes at pH = 3.50 ; $H_2O$; $T$ = 25.0°C; $[Ag^{+}]_{init}$ = 5.10$^{-3}$ M, $[L_4]_{init}$ = 7.5 x 10$^{-3}$ M.

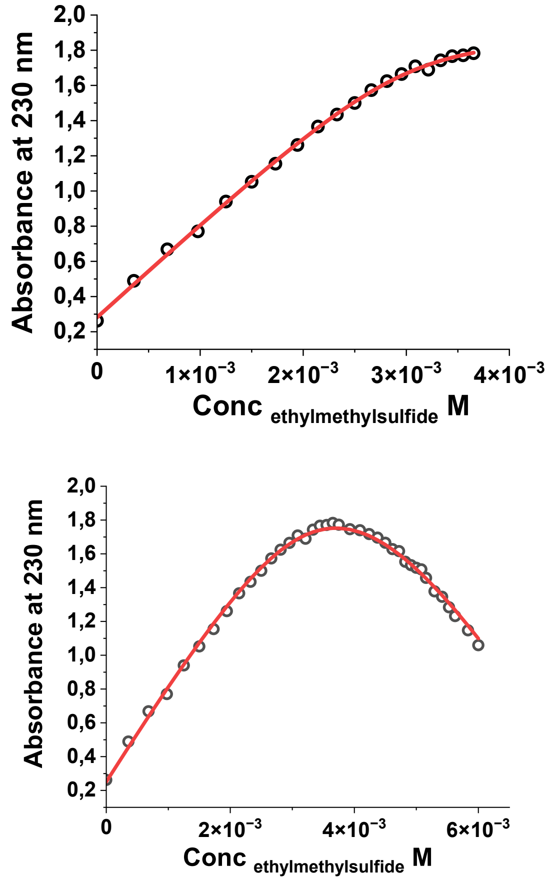

**Figure S11.** Absorbance at 230 nm as a function of the **ethylmethylsulfide** concentration. $H_2O$; $T$ = 25.0°C; $[Ag^{+}]_{init}$ = 5.10$^{-3}$ M, $[L_4]_{init}$ = 7.5 x 10$^{-3}$ M .

### 2. Computational Methods

#### 2.1 Calculation of the target silver(I)–toluene distance

We performed a relaxed scan in vacuum at the wB97XD/Def2TZVP level of theory using the program Gaussian 16.[14–16] Therefore, a Z-matrix was defined where the distances between the silver(I) ion and the six carbon atoms of the toluene ring were described by a single variable CAAG. This ensures equal distances between the silver(I) ion and the aromatic carbon atoms during the scan. Then a relaxed scan along the distance CAAG was performed. Finally, the potential energy profile was determined for the distance between the silver(I) ion and the center of the ring (by projecting the distance CAAG on the normal of the ring running through its center). The chosen target distance of 2.3 Å (as reported in Table 2 of the main text) was the minimum of this potential energy profile.

#### 2.2 Calculation of the binding constant of the sidechain fragments

##### 2.2.1 Setup of systems and software for all-atom simulations

The solvated structure of each sidechain fragment was obtained from the corresponding full amino acid residue and converting the $C^{\beta}$ atom into to a standard methyl group using the CHARMM software.[17] Therefore, the carbon atom of the terminal methyl-group was set to -0.27 *e*. The charge of all aliphatic hydrogens was set to 0.09 *e*. The CHARMM atom type was changed to CT3 and HA3 for the corresponding carbon and hydrogen atoms, respectively, of the methyl group. In the case of histidine (Hse tautomer), the conversion of the $C^{\beta}$ atom was problematic because its partial charge is not in line with a standard, charge-neutral methylene group and bonded force-field parameters with the imidazole ring are missing. Therefore, we decided to convert the $C^{\alpha}$ atom instead which avoided these difficulties.

CHARMM36m force field was used to model the sidechain fragments and either cTIP3P or sTIP3P water molecules were placed in a cubic box of side equal to 30 Å.[18,19] As a result, two different sets of simulations were defined according to the water type. One silver(I) ion was placed in each box, without further ions.

VMD v1.9.4a57 was used to visualize the trajectories.[20] All pictures of the structures were taken with UCSF Chimera v.1.17.1.[21]

##### 2.2.2 Umbrella sampling (US) of fragments

*General:* US simulations were performed with OpenMM 7.7 on a single NVIDIA GPU (RTX2080 or similar).[22] Langevin integrator was used to keep the temperature constant at 298.15 K in all the simulations, with a friction coefficient of 1/ps and a timestep of 1 fs for equilibration runs and 2 fs for production runs with SHAKE constraints for covalent hydrogen bonds.[23] Monte Carlo barostat was used to keep the pressure at 1 bar during production runs with a coupling frequency of 0.2 ps.[24,25] Long-range electrostatic interactions were treated by Particle Mesh Ewald scheme (PME) with an error tolerance of 0.0005.[26] The LJ interactions were switched to zero between 10 and 12 Å.

*US setup & simulations*: The system was biased along the reaction coordinate defined by the distance between the silver(I) ion and the interacting atom of the fragment (binding distance). The OpenMM CustomCentroidBondForce feature was used to define harmonic biasing potentials in 16 or 17 windows (depending on the system) to guarantee a partial superposition of the distance distributions between them. The force constants for the biasing potentials were chosen taking into account the experimentally measured binding affinities. Centers of the harmonic potentials ranged between 2 and 15 Å. For each window, an equilibration of 100 ps in the NVT ensemble was followed by four blocks of production runs of 10 ns each in the NPT ensemble. As a result, a total simulation time of 640 ns (680 ns with 17 windows) for each sidechain fragment was performed.

##### 2.2.3 PMF profile and structures of propanoate

Here is presented the PMF profile of propanoate in cTIP3P and sTIP3P water models. Structures along the PMF are reported for the cTIP3P model. The black line indicates the profile with HFE parameters only, and no red line is present because no parameterization was performed.

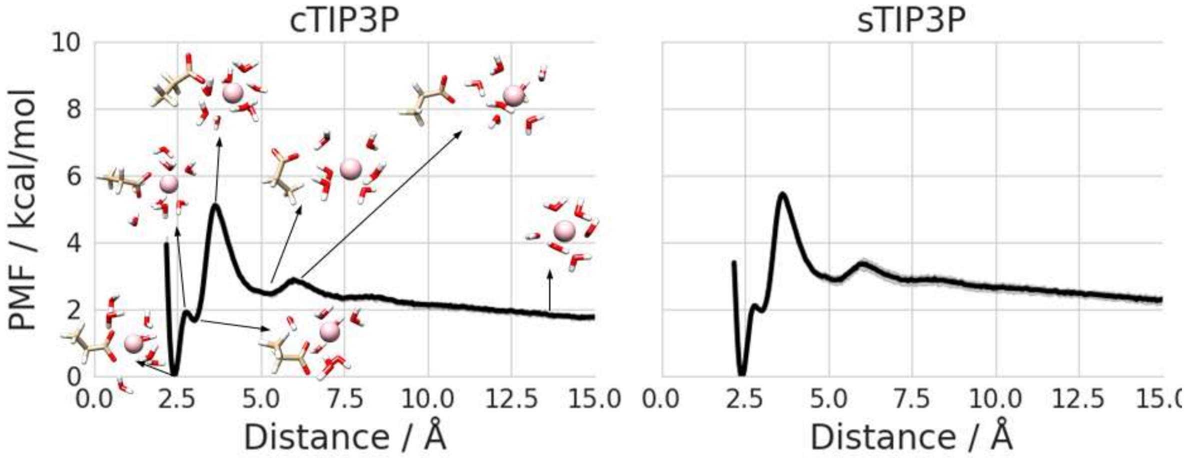

**Figure S12.** PMF profiles and structures of propanoate with HFE only parameters (black line) in cTIP3P (left) and sTIP3P (right) water models. PMF curves are plotted together with the standard error of the mean obtained from an average over four blocks (lighter colors, barely visible due to the small magnitude). For propanoate, the distance refers to the one from the carboxylic C atom. Structures are reported for the cTIP3P water model.

##### 2.2.4 Determination of the binding constant

For the calculation of the silver(I) binding constant of the sidechain fragments, the PMF as a function of the binding distance was calculated with WHAM for each block.[27] On this purpose, 260 bins and an error tolerance of 0.000001 were adopted. From each resulting PMF profile, the binding constant was calculated as indicated in equation 13 of Ref. [28] which reduces in our case to:

$$K_{bind,calc} = \frac{S^* \, I^*}{1661} \, \text{Å}^{-3} M^{-1} \tag{S1}$$

where:

$$S^* = 4\pi(r^*)^2 \tag{S2}$$

$$I^* = \int_{site} dr \, e^{-\beta[\mathcal{W}(r)-\mathcal{W}(r^*)]} \tag{S3}$$

and $\boldsymbol{r}^*$ is the distance when silver(I) ions are completely unbound (about 14 Å), $\mathcal{W}(\boldsymbol{r})$ is the PMF as a function of the binding distance $\boldsymbol{r}$; $\mathcal{W}(\boldsymbol{r}^*)$ is the PMF at the distance $\boldsymbol{r}^*$ in the bulk. The integration is over the binding site ($\boldsymbol{r} < 5$ Å). In practice, we used the trapezoidal rule to integrate Equation S3.

The final value of $K_{bind,calc}$ was obtained by the mean value over the four blocks. In addition, the standard error of the mean is reported in Table 2 (main text) for each sidechain fragment.

#### 2.3 Calculation of the binding constant of the tetrapeptides

##### 2.3.1 Setup of tetrapeptides systems and software for all-atom simulations

The structures of each tetrapeptide were obtained with the Build feature of Pymol to generate an initial .pdb file.[29] No cappings were added at this stage. The molecule was then loaded on the CHARMM-GUI webserver to obtain input files for OpenMM.[30,31] An acetyl moiety (ACE) and amide (CT2) moiety were used as termini cappings as in the experiments, and the protonation state of histidine was explicitly set to Hse. Waters and ions were set as for the sidechain fragments.

**Table S3:** Timings for all the scaled-potential MD simulations.

| System | Water model | Number of Ag(I) ion | Number of replicas | Time per replica (μs) | Total time (μs) |
|---|---|---|---|---|---|
| HEFM | cTIP3P | 1 | 8 | 0.8 | 6.4 |
| | sTIP3P | 1 | 8 | 0.8 | 6.4 |
| MNEH | cTIP3P | 1 | 8 | 0.8 | 6.4 |
| | sTIP3P | 1 | 8 | 0.8 | 6.4 |
| HAAM | cTIP3P | 1 | 8 | 0.8 | 6.4 |
| | sTIP3P | 1 | 8 | 0.8 | 6.4 |
| MAAH | cTIP3P | 1 | 8 | 0.8 | 6.4 |
| | sTIP3P | 1 | 8 | 0.8 | 6.4 |
| LP1 | cTIP3P | 0 | 8 | 5.0 | 40.0 |
| | sTIP3P | 0 | 8 | 5.0 | 40.0 |
| | cTIP3P | 1 | 8 | 5.0 | 40.0 |
| | sTIP3P | 1 | 8 | 5.0 | 40.0 |
| | cTIP3P | 5 | 8 | 2.5 | 20.0 |
| | sTIP3P | 5 | 8 | 1.0 | 8.0 |

##### 2.3.2 Scaled-potential Molecular Dynamics (MD) simulations of (tetra)peptides

The scaled-potential MD simulations were performed with OpenMM 7.7 on one NVIDIA GPU.[22,32] Unless explicitly specified, the same simulation settings were used as in the case of the US simulations. Due to the very high values of the experimental binding constants of the tetrapeptides, the $\varepsilon^{NBFIX}$ parameters were scaled down to ~40% for the interaction between silver(I) and the atom $N^{\delta}$ of histidine (atom type NR2; $\varepsilon'^{NBFIX}$ = -1.88 kcal/mol) and to ~50% for atom S of methionine (atom type S; $\varepsilon'^{NBFIX}$ = -7.20 kcal/mol) in order to accelerate the dissociation. Both (reversible) association and dissociation are required for the proper calculation of the binding constant. Due to the inter-dependency of $\varepsilon^{NBFIX}$ and $R^{NBFIX}$, the latter was modified accordingly in order to keep the same equilibrium distance.

Eight replicas were created by modifying the initial coordinates file, where the silver(I) ion was placed in each corner of the cubic box. For each replica, 10000 steps of minimization with a force tolerance of 23.9 kcal/mol were followed by an equilibration in the NVT ensemble of 100 ps.

Production runs in the NPT ensemble were performed for 800 ns, for a total production time of 6.4 $\mu s$ for each tetrapeptide. The simulations were repeated for both cTIP3P and sTIP3P. In the case of the LP1 peptide, in addition to the simulations with silver(I) as described above, eight replicas of simulations without the ion were performed in both cTIP3P and sTIP3P. As a result, four cases were defined combining the presence/absence of the silver(I) ion with the two water models.

Table S3 presents a synthesis of the production runs.

##### 2.3.3 Determination of the binding constants

Hamiltonian reweighting was adopted to retrieve the correct binding constants of the (tetra)peptides from the scaled-potential MD simulations by first calculating the bias $b_k$ of the $k$-th snapshot of the simulation trajectory from the sum of the corrections of the $i$-th and the $j$-th atoms, and then deriving its weight $w_i$:[33]

$$b_k = \sum_{i,j}^{atoms} \varepsilon'^{NBFIX}_{ij} \left[ \left( \frac{R'^{NBFIX}_{ij}}{r_{ij}} \right)^{12} - 2 \left( \frac{R'^{NBFIX}_{ij}}{r_{ij}} \right)^{6} \right] - \varepsilon^{NBFIX}_{ij} \left[ \left( \frac{R^{NBFIX}_{ij}}{r_{ij}} \right)^{12} - 2 \left( \frac{R^{NBFIX}_{ij}}{r_{ij}} \right)^{6} \right] \tag{S4}$$

$$w_k = \exp(-b_k / k_B T) \tag{S5}$$

where, $i$ is the atom type of silver(I) and $j$ is either $N^{\delta}$ (histidine), S (methionine), $C_{aromatic}$ (phenylalanine). $R^{NBFIX}_{ij}$ and $\varepsilon^{NBFIX}_{ij}$ are the NBFIX parameters for the interaction pair $ij$. The scaled NBFIX parameters (as used in the scaled-potential MD simulations) are indicated with a prime sign. Since $\varepsilon'^{NBFIX}_{ij} < \varepsilon^{NBFIX}_{ij}$ it is $b_k \leq 0$. As a consequence, $w_k \geq 1$ with the special case $w_k = 1$ if $b_k = 0$, that is when silver(I) and the peptide are not interacting. Note that Eqns. S4 & S5 are slightly incorrect for large values of $r_{ij}$ since we used a force-switching scheme of the Lennard-Jones interactions (including NBFIX corrections) at large values of $r_ij$. The error can, however, be neglected; we verified this for the tetrapeptide with the highest affinity (HEFM).

The binding constant of two monomers were derived from counting the number of states where the two monomers are associated ($n_1$) and where are dissociated ($n_0$):[34]

$$K_{bind,calc} = N_{Av} \frac{n_1}{n_0} (v - v_D) \tag{S6}$$

where $N_{Av} = 6.022 \ 10^{23}$ mol$^{-1}$ is Avogadro's number; $n_1/n_0$ is the ratio of the counts of bound and unbound states; $v$ and $v_D$ are namely the average volume of the simulation and the dimer volume, in liter. For small systems such as a tetrapeptide we can assume that the dimer volume is neglectable compared to the simulation volume, that is: $v_D \ll v$. To verify this assumption, we tested two different simulation volume sizes ($30^3$ and $50^3$ Å$^3$).

In the context of Hamiltonian reweighting, the ratio between the number of bound and unbound states became the ratio between the weights associated to those states. Once the weights for each frame of the trajectory were obtained, the binding constant for the four tetrapeptides was calculated with the following formula:

$$K_{bind,calc} \cong \quad cv\frac{\sum^{bound} w_i}{\sum^{unbound} w_i} \tag{S7}$$

where $c$ = 1/1661 $Å^{-3}M^{-1}$ is a conversion factor, $v$ is the average volume of the simulation box in $Å^3$. In the case where no bias is applied, all the weights are equal to 1 and Equation S7 boils down to Equation S6. To determine whether a state was "bound" or "unbound", we evaluated the minimum distance between any atom of each (tetra)peptide and silver(I). If the value was within a threshold distance of 5 Å, the weight $w_i$was summed up to the numerator, otherwise to the denominator. *Ad-hoc* Python scripts were developed for this purpose with the aid of MDAnalysis package.[35,36]

The $K_{bind,calc}$ was determined for each replica and the mean value as well as the standard error of the mean are reported in Table 3 (main text) for each tetrapeptide.

##### 2.4 Calculation of the α-helical content of LP1

The LP1 oligopeptide was setup and simulated in analogy to the tetrapeptides (Section 2.3.1) using eight replicas with 0, 1 or 5 silver(I) ion(s). To probe the $\alpha$-helical content of LP1, the ALPHARMSD function of PLUMED was applied *a posteriori* on the trajectory.[37,38] In particular, the function can be applied on a stretch of six subsequent residues with a score between 0 (no $\alpha$-helix content) and 1 (optimal $\alpha$-helix). Since it is not known experimentally which residues of LP1 fold into a helix, we selected for the analysis all 14 residues. In practice, the ALPHARMSD function was determined for each stretch of six subsequent residues; for LP1 there are in total 9 such stretches. For each snapshot the sum of the 9 ALPHARMSD values was calculated and divided by 9 to obtain a normalized $\alpha$-helical score. Using the statistical weights from the Hamiltonian reweighting an average of this $\alpha$-helical score was performed using snapshots every 500 ns on the full trajectory for each replica. After that, the weighted scores of the eight replicas were averaged, and the error bars obtained as standard errors of the mean.
